# Constitutive nuclear accumulation of endogenous alpha-synuclein in mice causes motor dysfunction and cortical atrophy, independent of protein aggregation

**DOI:** 10.1101/2021.10.13.464123

**Authors:** Haley M. Geertsma, Terry R. Suk, Konrad M. Ricke, Kyra Horsthuis, Jean-Louis A. Parmasad, Zoe Fisk, Steve M. Callaghan, Maxime W.C. Rousseaux

**Affiliations:** Department of Cellular and Molecular Medicine, University of Ottawa, Ottawa, ON, K1H8M5; University of Ottawa Brain and Mind Research Institute, Ottawa, ON, K1H8M5; Ottawa Institute of Systems Biology, Ottawa, ON, K1H8M5; Eric Poulin Center for Neuromuscular Diseases, Ottawa, ON, K1H8M5

**Keywords:** Nuclear alpha-synuclein, Parkinson’s disease, Neurodegeneration, Darpp-32

## Abstract

**Background:** A growing body of evidence suggests that nuclear alpha-synuclein (αSyn) plays a role in the pathogenesis of Parkinson’s disease (PD). However, this question has been difficult to address as controlling the localization of αSyn in experimental systems often requires protein overexpression, which affects its aggregation propensity.

**Methods:** We engineered *Snca*^*NLS*^ mice which localize endogenous αSyn to the nucleus. We characterized these mice on a behavioral, histological, and biochemical level to determine whether the increase of nuclear αSyn is sufficient to elicit PD-like phenotypes.

**Results:** *Snca*^*NLS*^ mice exhibit age-dependent motor deficits and altered gastrointestinal function. We found that these phenotypes were not linked to αSyn aggregation or phosphorylation. Through histological analyses, we observed motor cortex atrophy in the absence of midbrain dopaminergic neurodegeneration. We sampled cortical proteomes of *Snca*^*NLS*^ mice and controls to determine the molecular underpinnings of these pathologies. Interestingly, we found several dysregulated proteins involved in dopaminergic signaling, namely Darpp-32, which we further confirmed was decreased in cortical samples of the *Snca*^*NLS*^ mice compared to controls via immunoblotting.

**Conclusions:** These results suggest that chronic *endogenous* nuclear αSyn can elicit toxic phenotypes in mice, independent of its aggregation. This model raises key questions related to the mechanism of αSyn toxicity in PD and provides a new model to study an underappreciated aspect of PD pathogenesis.

## Background

Alpha synuclein (αSyn) is a protein notorious for its involvement in Parkinson’s disease (PD) pathogenesis. For one, it is a primary constituent of Lewy bodies and Lewy neurites, pathological hallmarks of PD (1). Moreover, copy number variations and missense mutations in the αSyn gene, *SNCA*, cause genetic forms of PD, further reinforcing its involvement in disease etiology (2–8). αSyn was first described as a presynaptic and nuclear protein (9). However, nuclear αSyn has largely been overshadowed by a focus on its cytoplasmic form, likely due to the cytoplasmic localization of Lewy bodies. Despite this, several studies have linked nuclear αSyn to PD on multiple levels: in cell (10–14) and animal (6,15–18) models of PD, as well as in brain tissue from individuals with αSyn pathologies (synucleinopathy). These studies examined the role of nuclear αSyn by overexpressing it together with mutations or a nuclear localization signal (NLS) (12,15,17–20), or upon toxin exposure (16), hinting at a role for nuclear αSyn in disease pathogenesis by its involvement in DNA binding (21,22) or histone modification (17) to alter transcription, and in DNA repair (23). While these studies support a link between nuclear αSyn and PD, its specific role in disease – whether deleterious or beneficial – remains clouded due to the reliance on its overexpression or exogenous stressors, making it difficult to parse out the driver of toxicity in the absence of αSyn aggregation.

To directly test the consequence of chronic αSyn mislocalization to the nucleus *in vivo*, without resorting to protein overexpression, we engineered a mouse model to endogenously express αSyn with a C-terminal NLS-Flag tag driving its mislocalization to the nucleus. We extensively characterized these mice at the behavioral, histological, and biochemical level to assess whether chronic nuclear localization of αSyn causes age-dependent phenotypes resembling PD or related αSyn proteinopathies.

## Methods

### Mouse Design and Engineering

#### Mouse Engineering

To generate the *Snca*^*NLS*^ mice, Cas9 protein was complexed to a sgRNA targeting the 3’ locus of *Snca* (sgRNA target sequence: 5’-TTGGTAGCCTTCCTAATATC-3’); and, together with a single-stranded oligodeoxynucleotide (ssODN) repair template (sequence: 5’-CACTGTGAAGCAGACAGTTGATATCTGTCACTTCACTGACAAGGCATGCTGTTATTATTTTCTTTTTCTGATATTAGGAAGGCTACCAAGACTATGAGCCTGAAGCCGACTACAAGGACGACGACGACAAGTAAGAATGTCATTGCACCCAATCTCCTAAGATCTGCCGGCTGCTCTTCCATGGCGTACAAGTGCTCAGT -3’; IDT Ultramer), were injected and implanted into pseudo-pregnant FVB female mice (24). Three founder mice were generated and backcrossed onto a ∼99.75% pure C57Bl6/J background (Taconic 1450 SNP analysis, sequencing and subsequent backcrossing; see **Supplementary Figure 1A, B**), before being expanded onto a mixed C57Bl6/J;C57Bl6/NCrl background. One line was selected for subsequent extensive characterization and will soon be made available through the Jackson Laboratory (Jax Stock No. 036763).

#### Genotyping

A small (<1 mm) tail sample is digested prior to PCR amplification using primers outside of the sequence covered by the ssODN used for the initial mouse line. Forward: 5’-TTTTATCTGATTGAAATGATGAGC-3’; Reverse: 5’-ATGACTGGGCACATTGGAA-3’. PCR protocol: 95°C for 2 minutes, (95 °C for 30 seconds, 56 °C for 30 seconds, 72 °C for 30 seconds) repeated for 35 cycles, 72 °C for 5 minutes. Mutant allele: 273 bp; Wildtype allele: 225 bp.

### Mouse Husbandry

All mice were housed with up to 5 mice per cage on a 12-hour light-dark cycle. Mice were fed *ad libitum* and all husbandry was performed by the uOttawa Animal Care and Veterinary Services staff. All animal work was done under the approved breeding (CMMb-3009 and CMMb-3654) and behaviour (CMMe-3091) protocols approved under the uOttawa Animal Care Committee. All mice were handled daily for 1 week prior to all behavior testing and both male and female mice were used in all experiments.

### Behavior

#### Open Field

Lighting in the behavior room was set to 100 lux and mice were habituated for 60 minutes prior to testing. Mice were placed into the open field box (45 cm^3^) for 10 minutes with their movement recorded/analyzed with Ethovision software (Noldus Information Technology).

#### Fecal Pellet Output

Upon completion of the Open Field test, the number of fecal pellets excreted during the 10-minute trial were quantified.

#### Beam Break

Single mice were placed into a clean cage with access to food and water *ad libitum* for 24 hours at the standard 12-hour light-dark cycle with their movement recorded/analyzed via Fusion software.

#### Nesting

Directly following Beam Break testing, one square of nestlet (5 cm^2^ cotton pad) was placed in each Beam Break cage for 17-19 hours. Following this, the nestlets were scored on a scale of 1-5 as described in Deacon 2006 (25), with 1 and 5 representing minimal and maximal nest quality, respectively. *Y maze Forced Alternation*: Mice were provided with extra-maze (irregular black cue with squared edges on right wall, black triangle on left wall) and intra-maze (Arm 1 has solid black rectangle, Arm 2 has horizontal bars, Arm 3 has diagonal stripes) cues. The room was set to 60 lux and mice were habituated for 60 minutes prior to testing. During the first 5-minute trial, Arms 2 and 3 were alternatively. Following the 30-minutes inter-trial interval, the mice were placed back into the Y maze apparatus for 5 minutes without any blocked arms and their movements were recorded/analyzed with EthoVision software.

#### Adhesive Removal

After a 60-minute habituation, the home cage of the mice was lightly wiped to remove all bedding material. The mice were individually placed back into the emptied home cage for a 1-minute habituation. Next, a 1 cm^2^ square of medical adhesive was placed on each forepaw and the mice were placed back into the wiped home cage where the time to remove adhesive was measured up to a 2-minute maximum.

#### Pole test

Following a 60-minute habituation, mice were placed on a textured metal pole (8 mm diameter, 55 cm tall) ∼3 cm from the top facing upwards. The mice were given up to 1 minute to turn around (facing downwards) and up to 1 minute to descend the pole.

#### Rotarod

Following a 60-minute habituation, mice were placed on a rod (IITC Life Sciences) rotating from 4-40 revolutions per minute over 5 minutes for 4 trials per day with a 10-minute inter-trial interval. This was repeated for 3 days total.

#### DigiGait

Following a 60-minute habituation, mice were placed in the DigiGait treadmill (Mouse Specifics Inc.). The treadmill ran at 22 cm/s (3- and 9-months timepoints) or 18 cm/s (18-month timepoint) with a 0° incline. 3 seconds of continuous movement was recorded using DigiGait Imager software and was then analyzed with DigiGait Analysis software.

#### Fear Conditioning

NaÏve age- and sex-matched mice were used to obtain the optimal intensity of foot shock. On day 1 of testing, mice were placed into the Fear Conditioning apparatus (Noldus Information Technology) for 6 minutes during which time the mice experienced 3 tone-shock pairings (30-second tone co-terminated with a 2-second foot shock). On day 2, mice were placed into the same Fear Conditioning apparatus for 6 minutes with no tone or foot shock. On day 3, mice were placed into a different Fear Conditioning apparatus, now altered into a triangular shape with solid floor and a vanilla scent for 3 minutes with no tone then 3 minutes with the same 30-second tone but no foot shock. Freezing was measured/analyzed with EthoVision on all 3 testing days.

#### Fecal Pellet Composition

Mice were placed in a clean cage and their fecal pellets were collected over a 1-hour period. These pellets were weighed (wet weight) then desiccated at 65 °C for 19 hours and reweighed (dry weight). The differences were calculated between these values to determine the water content.

### Histology

See **Supplementary Table 1** for a comprehensive list of antibodies used in this study.

#### Perfusion

Mice were sedated with 120 mg/kg Euthanyl (DIN00141704) then perfused with 1X phosphate buffered saline (PBS) then 4 % paraformaldehyde (PFA). Brains were then extracted and incubated in 4 % PFA for 72 hours prior to a 3-step sucrose dehydration with 10, 20 and 30% sucrose (24 hours each). Next, brains were flash frozen for 1 minute in -40 °C isopentane and cryosectioned at 40 μm. *Immunofluorescent staining:* Cryosectioned tissue was mounted on a slide then blocked with blocking buffer (0.1 % Triton X-100, 10 % normal horse serum in 1X PBS) then incubated in primary antibody overnight at 4 °C. Next, the sections were incubated in secondary antibody before drying at room temperature for 2 minutes. The sections were then covered with #1.5 coverslips and Vectashield Antifade Mounting Medium with DAPI (MJS Biolynx cat# VECTH1200).

#### Quick decapitation, fixation

Mice were euthanized by isoflurane inhalation followed by decapitation and brains were quickly extracted. Brains were then submerged in 10 % buffered formalin for 72 hours prior to paraffin embedding and sectioning at 5 μm.

#### Diaminobenzidine (DAB) staining

Paraffin-embedded sections were deparaffinized in serial baths of xylenes and ethanol prior to a sodium citrate (10 mM sodium citrate, 0.05 % Tween-20, pH 6) antigen retrieval (20 minutes at 95 °C) and 0.9 % H_2_O_2_ treatment (10 minutes). Next, sections were blocked in blocking buffer (0.1 % Triton X-100, 10 % normal horse serum in 1X PBS) then incubated in primary antibody overnight at 4 °C. The following day the sections were incubated in secondary and tertiary antibody solution before exposure to DAB, dehydrating in baths of ethanol and xylenes, and covering the tissue with Permount (Fisher Scientific cat# SP15-100) and #1.5 coverslips.

#### Toluidine Blue and H&E staining

Staining was performed by the Louise Pelletier Histology Core facility at the University of Ottawa on paraffin-embedded 5 μm sectioned mouse brain tissue using the Leica Autostainer XL. Briefly, the sections were deparaffinized and exposed to Toluidine Blue for 10 minutes.

For H&E, sections were deparaffinized, exposed to hematoxylin for 7 minutes and eosin for 30 seconds. Then, sections were dehydrated and covered with #1.5 coverslips.

#### Stereology

Stereology was performed as previously described (26). Briefly, for each mouse, 8 cryosections were stained for tyrosine hydroxylase (TH) and quantified using StereoInvestigator software (version 11.06.2). The sections (40 µm) began at the outer limit of the substantia nigra (SNc) and every 6^th^ section was used (Bregma -2.54 mm to -3.88 mm). Mean section thickness was determined during counting at a frequency of 10 frames (roughly 3 measurements per hemisphere). The SNc was sampled by randomly translating a grid with 150 μm × 150 μm squares in the outlined SNc and applying an optical fractionator consisting of a 75 μm × 75 μm square.

### Biochemistry

See **Supplementary Table 1** for a comprehensive list of antibodies used in this study.

#### Serial Extraction

A 1 mm^3^ punch of cortical brain tissue from 18-month-old mice was homogenized and resuspended in a series of increasingly stringent buffers beginning with 100 µL of TSS Buffer (140 mM NaCl, 5 mM Tris-HCl), then 100 µL TXS Buffer (140 mM NaCl, 5 mM Tris-HCl, 0.5% Triton X-100), then 100 μL SDS Buffer (140 mM NaCl, 5 mM Tris-HCl, 1% SDS), as previously described (27). Total protein levels were measured using the Pierce^™^ BCA Assay Kit (Thermo Fisher cat# 23225).

#### Western Blot

Protein samples were loaded into a 12 % polyacrylamide gel and subsequently transferred to a 0.2 µm nitrocellulose membrane. Membranes were blocked in a 5 % milk solution then incubated in primary antibody (diluted in 2 % bovine serum albumin) overnight at 4 °C. Next, the membrane was incubated in a horseradish peroxidase-conjugated secondary antibody diluted in 5% milk solution. Then, the membrane was rinsed with ECL Clarity solution (Bio-Rad cat# 1705061) and imaged with the GE ImageQuant LAS 4000.

#### RNA Extraction and Real-Time Quantitative PCR

RNA was extracted from mouse brain homogenate using Trizol-Chloroform extraction (Invitrogen^™^ User Guide: TRIzol Reagent version B.0). Briefly, mouse brains were homogenized in 3 mL of PEPI Buffer (5 mM EDTA, 1X protease inhibitor (GenDEPOT cat# P3100-020), in 1X PBS) using a dounce homogenizer. 3 % of homogenate was added to 1 mL of TRIzol Reagent (Fisher Scientific cat# 15-596-026) and RNA was isolated following the user guide referenced above. cDNA was synthesized using 5X All-in-One RT Master Mix Kit (Bio Basic cat# HRT025-10). Real-time quantitative PCR was performed using Green-2-Go qPCR Master Mix (Bio Basic cat# QPCR004-S) with 25 ng cDNA per reaction and primers targeting mouse *Gapdh* (Forward: 5’-GGAGAGTGTTTCCTCGTCCC-3’, Reverse: 5’-ATGAAGGGGTCGTTGATGGC-3’), *Hprt1* (Forward: 5’-TGATAGATCCATTCCTATGACTGTAGA-3’, Reverse: 5’-AAGACATTCTTTCCAGTTAAAGTTGAG-3’), and *Snca* (Forward: 5’-GAAGACAGTGGAGGGAGCTG-3’, Reverse: 5’-CAGGCATGTCTTCCAGGATT-3’). Reactions were run on BioRad CFX96 thermocycler (protocol: 95 °C for 5 minutes, 40 cycles of 95 °C for 15 seconds and 60 °C for 60 seconds, then melting curve). *Snca* Ct values were standardized to the average of *Hprt1* and *Gapdh*.

#### Dopamine and Metabolite Measurement via Liquid Chromatography-Mass Spectrometry/Mass Spectrometry (LC-MS/MS)

Striatal punches (2 mm i.d., 3 mm thick section) were extracted from 18-month mouse brains and weighed prior to submitting to The Metabolomics Innovation Centre (TMIC). 50 µL of tissue extraction buffer was added to each sample tube followed by homogenization and centrifugation. Supernatant was used for LC-MS/MS analysis to get the above concentrations in the unit of µM. TMIC staff applied a targeted quantitative metabolomics approach to analyze the samples using a reverse-phase LC-MS/MS custom assay. This custom assay, in combination with an ABSciex 4000 QTrap (Applied Biosystems/MDS Sciex) mass spectrometer, can be used for the targeted identification and quantification of measure dopamine (DA), homovanillic acid (HVA), 5-hydroxyindoleacetic acid (5-HIAA), and 3,4-dihydroxyphenylacetic acid (DOPAC). The method combines the derivatization and extraction of analytes, and the selective mass-spectrometric detection using multiple reaction monitoring (MRM) pairs. Isotope-labeled internal standards and other internal standards are used for metabolite quantification. The custom assay contains a 96 deep-well plate with a filter plate attached with sealing tape, and reagents and solvents used to prepare the plate assay. First 14 wells were used for one blank, three zero samples, seven standards and three quality control samples. For all metabolites except organic acid, samples were thawed on ice and were vortexed and centrifuged at 13,000x *g*. 10 µL of each sample was loaded onto the center of the filter on the upper 96-well plate and dried in a stream of nitrogen. Subsequently, phenyl-isothiocyanate was added for derivatization. After incubation, the filter spots were dried again using an evaporator. Extraction of the metabolites was then achieved by adding 300 µL of extraction solvent. The extracts were obtained by centrifugation into the lower 96-deep well plate, followed by a dilution step with MS running solvent.

For organic acid analysis, 150 µL of ice-cold methanol and 10 µL of isotope-labeled internal standard mixture was added to 50 µL of sample for overnight protein precipitation. Then it was centrifuged at 13000x *g* for 20 min. 50 µL of supernatant was loaded into the center of wells of a 96-deep well plate, followed by the addition of 3-nitrophenylhydrazine (NPH) reagent. After incubation for 2h, BHT stabilizer and water were added before LC-MS injection.

Mass spectrometric analysis was performed on an ABSciex 4000 Qtrap® tandem mass spectrometry instrument (Applied Biosystems/MDS Analytical Technologies, Foster City, CA) equipped with an Agilent 1260 series UHPLC system (Agilent Technologies, Palo Alto, CA). The samples were delivered to the mass spectrometer by a LC method followed by a direct injection (DI) method. Data analysis was done using Analyst 1.6.2.

#### TMT10plex^*™*^ Proteomics via Liquid Chromatography-Mass Spectrometry (LC-MS)

Whole mouse cortex was dissected from 9-month-old mouse brain and peptides were isolated using the EasyPep^™^ Mini MS Sample Prep Kit (Thermofisher cat# A40006) following manufacturer instructions. These samples were labelled with the TMT10plex^™^ Isobaric Label Reagent Set (Thermofisher cat# 90406) then combined into a single tube and fractionated into 12 samples using the Pierce^™^ High pH Reversed-Phase Peptide Fractionation Kit (Thermofisher cat# 84868). Fractions 2, 3, 9, 10, 11, and 12 were combined due to low protein concentration (combined to have a consistent protein concentration with other fractions) and the 6 final fractions were submitted for LC-MS to the Ottawa Hospital Research Institute Proteomics Core Facility. LC-MS was performed using Orbitrap Fusion Lumos mass spectrometer with UltiMate 3000 RLSC nano HPLC (Thermo Scientific). Proteowizard MS-CONVERT was used to generate peak lists for preliminary qualitative analysis using MASCOT software version 2.7.0 (Matrix Science, UK). Protein identification and quantitative analysis was performed using MaxQuant (Tyanova, Nature Protocols 2016, 11:2301). The reference proteome for peptide spectrum matching was UniProt/Mus musculus (version 2020-10-06). The MaxQuant results were exported to Scaffold Q+S (Proteome Software, USA) for further analysis and viewing.

### Statistical Analyses

All statistical analyses were performed using GraphPad Prism (version 9.1.2) using the appropriate statistical test, either Student’s t-test for simple comparisons or One- or Two-way ANOVA followed by Bonferroni post-hoc analysis for multiple comparisons. The survival curve was analyzed using a log-rank Mantel-Cox test. The Mann-Whitney test with Benjamini-Hochberg correction was used with Scaffold (version 5.0.0) to analyze the TMT10plex^™^ mass spectrometry dataset. Statistical tests used, sample sizes, and p-values are delineated in each figure legend.

## Results

### Effective nuclear targeting of αSyn in *Snca*^*NLS*^ mice

To study if nuclear αSyn is sufficient to elicit age-related behavioral and pathological phenotypes, we generated a mouse line that targets endogenous αSyn to the nucleus via the knockin of an NLS-Flag tag on αSyn (*Snca*^*NLS*^). The NLS-Flag construct was targeted to the 3’ end of the *Snca* coding sequence with the modified *Snca-NLS-Flag* gene predicted to transcribe a fusion protein of wildtype αSyn with a C-terminal NLS-Flag tag (**Fig. 1A**). After generating and backcrossing mice (see Methods and **Supplementary Figure 1**), we confirmed the knockin via sequencing (**Supplementary Figure 1B**) and were able to distinguish between the genotypes via a PCR band shift (**Fig. 1B**) and a larger protein size via western blot (**Fig. 1C**). Mice were born at expected Mendelian ratios (**Supplementary Figure 1C**), confirming that insertion of this tag did not pose major developmental deficits.

**Figure 1:**
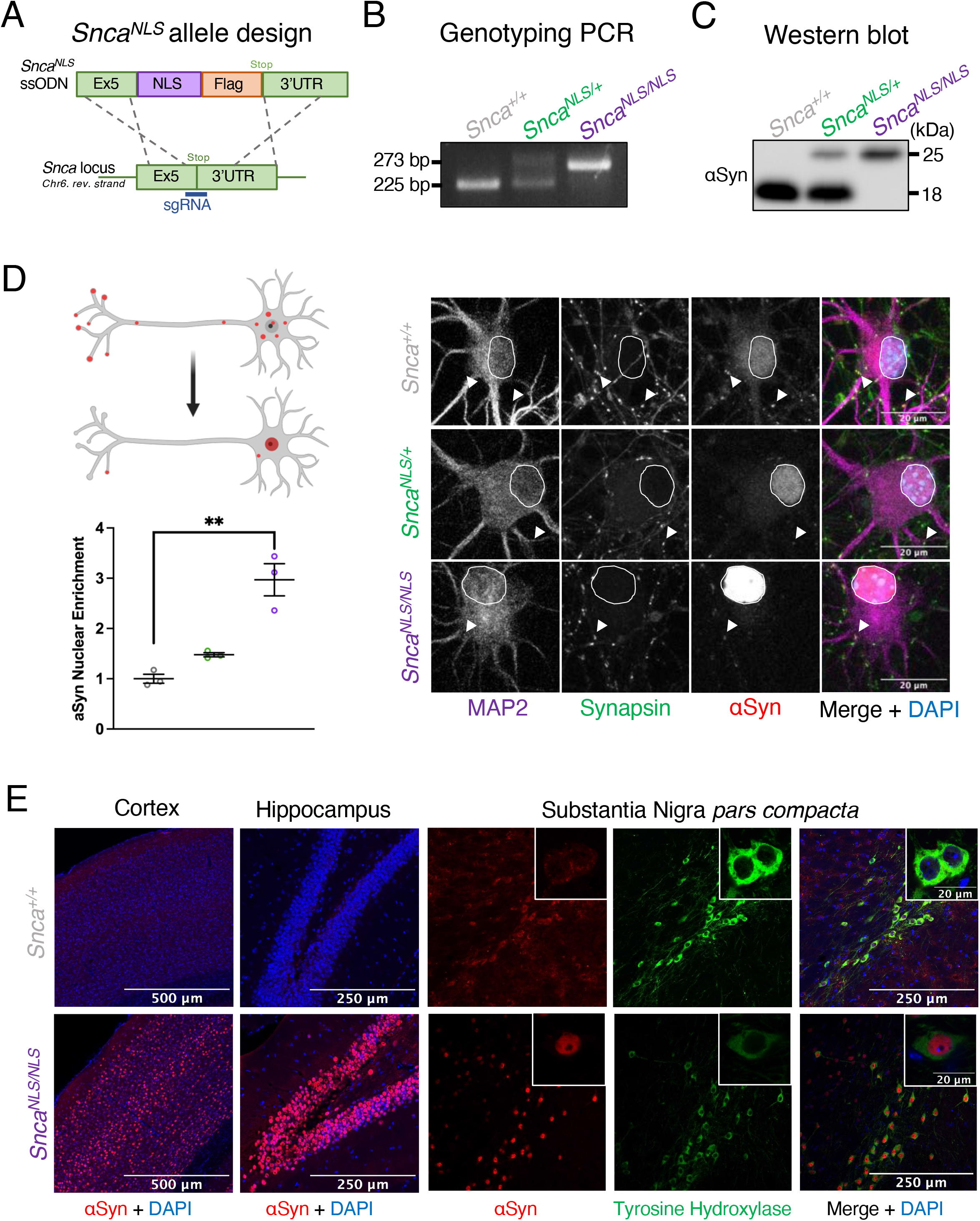
*Snca*^*NLS*^ mice effectively target αSyn to the nucleus in vitro and in vivo. A) *Snca*^*NLS-Flag*^ knock-in scheme with C-terminal NLS-Flag tag. Visualization of the knockin via (B) PCR and (C) western blot. D) Illustration of nuclear localization of αSyn (upper left) with protein quantification (lower left) of nuclear αSyn from primary cortical neurons in wildtype (top right panels), *Snca*^*NLS/+*^ (middle right panels), and *Snca*^*NLS/NLS*^ (bottom right panels) (n=3). White arrows denote presynaptic αSyn and nuclei circled in white E) Localization of αSyn in wild-type (top panels) and *Snca*^*NLS/NLS*^ (bottom panels) in the cortex, hippocampus, and SNc. One-way ANOVA with Bonferroni multiple comparison: ^**^ denotes p<0.01.

We next examined the efficiency of the NLS-Flag tag by quantifying the level of nuclear αSyn in primary cortical neuron cultures through immunofluorescent microscopy. We observed a 3-fold increase in nuclear αSyn in *Snca*^*NLS/NLS*^ and a 1.5-fold increase in *Snca*^*NLS/+*^ compared to *Snca*^*+/+*^ (wildtype) cells (**Fig. 1D**). This trend was consistent in stained adult mouse brain tissue (**Fig. 1E**). Importantly, this roughly corresponds to the 2.5-3-fold increase of nuclear αSyn which we and others have previously observed in post-mortem brain tissue from individuals with PD or other animal models of synucleinopathy, suggesting the model displays a disease-relevant increase of nuclear αSyn (6,15–18,26).

### Increased nuclear αSyn leads to an age-dependent motor decline, gastrointestinal dysmotility and premature lethality

To test whether chronic nuclear accumulation of αSyn is sufficient to elicit PD-like phenotypes over time, we subjected *Snca*^*NLS/NLS*^ mice and littermates to a battery of behavior tests at 3-, 9-, and 18-months of age. We found that the *Snca*^*NLS/+*^ and *Snca*^*NLS/NLS*^ mice performed similarly to wildtype at 3-months of age, with a mild motor deficit in the *Snca*^*NLS/NLS*^ mice appearing during beam break (**Fig. 2A**) and rotarod (**Fig. 2B**) tests. By 9-months, however, the *Snca*^*NLS/NLS*^ mice displayed a severe motor deficit in rotarod (**Fig. 2B**) as well as a delayed time to contact their forepaws in the adhesive removal test (**Fig. 2C**). Interestingly, *Snca*^*NLS/+*^ mice also exhibited a significant deficit on the rotarod test – suggestive of a dominant phenotype. Surprisingly, after aging these mice to 18 months, we observed milder motor phenotypes relative to wildtype controls; likely due to the 18-month wildtype mice showing increased difficulty at performing these tasks (**Supplementary Figure 2**).

**Figure 2:**
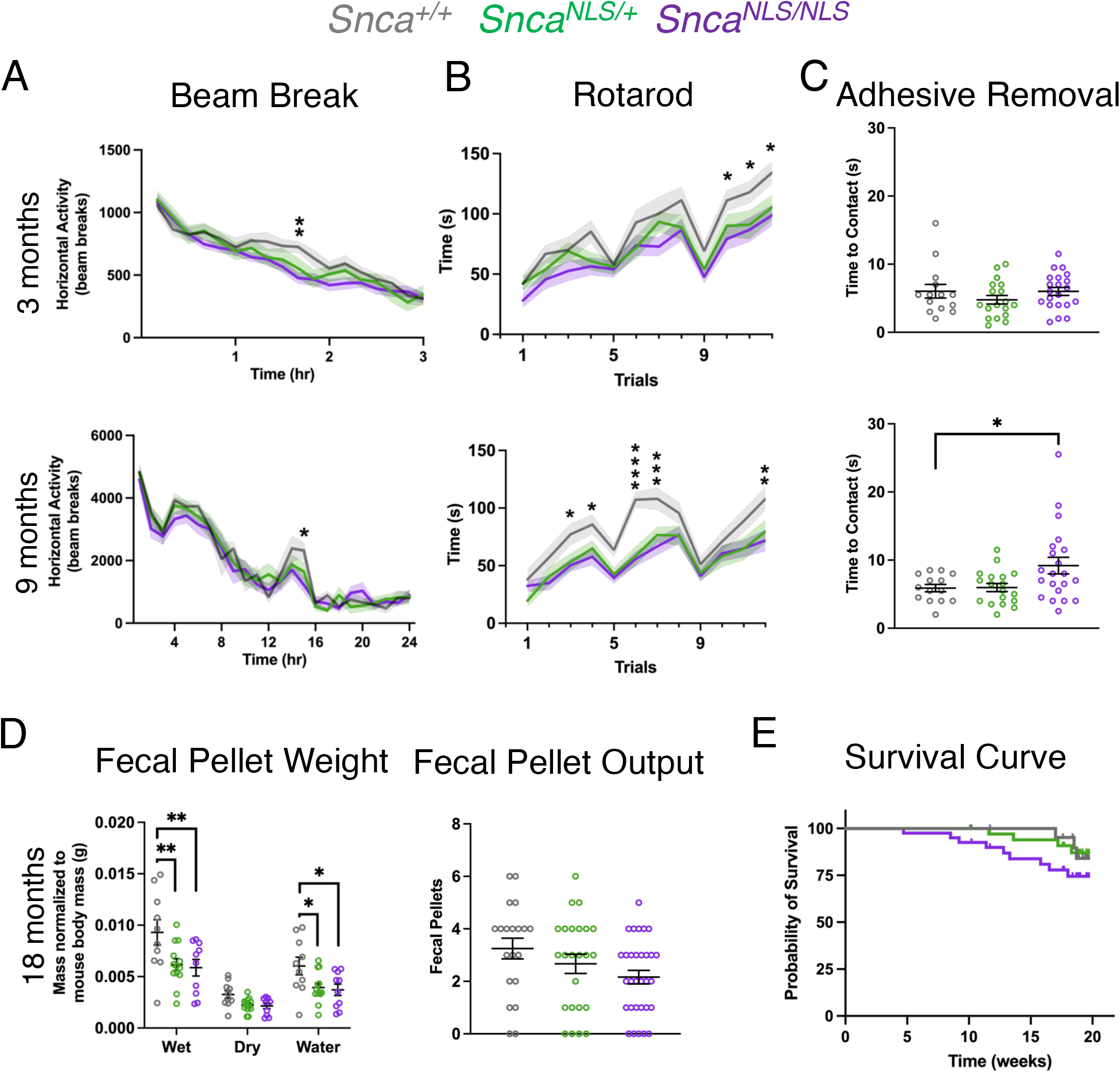
*Snca*^*NLS*^ mice exhibit significant motor and gastrointestinal dysfunction. Analysis of the 3-(top) and 9-month (bottom) mice for (A) Beam Break, (B) Rotarod, and (C) Adhesive Removal measuring time to contact their forepaws, (D) Fecal Pellet Weight measuring fecal weight and water content over 1 hour, and (E) Fecal Pellet Output measuring fecal pellets produced in 10 minutes, (n=10-21). F) Survival curve from all mice in the behaviour colony (n=32-58). One-Way (C,E) or Two-Way ANOVA (A,B,D) with Bonferroni multiple comparison or Log-rank (Mantel Cox) test (F): ns, ^*, **, ***^, and ^****^ denotes p>0.05, <0.05, <0.01, <0.001, and <0.0001, respectively.

With increasing awareness around non-motor symptoms in PD, we also measured cognition, anxiety, and overall wellness in the *Snca*^*NLS*^ line. We found that *Snca*^*NLS/NLS*^ and *Snca*^*NLS/+*^ mice performed similarly to their wildtype littermates in non-motor behavior tests at all timepoints (**Supplementary Figures 3-5**). In addition to motor decline, people living with PD often experience gastrointestinal difficulties such as constipation (28,29). To measure constipation in our mice, we examined fecal excretions in the span of an hour. We found that 18-month-old *Snca*^*NLS/NLS*^ mice excretions contained significantly less water than their wildtype counterparts (**Fig. 2D**) and a trend for decreased fecal matter production (**Fig. 2E**). Lastly, we observed a trend for early lethality in the *Snca*^*NLS/NLS*^ mice compared to their wildtype littermates, where 25% of *Snca*^*NLS/NLS*^ mice died by 20 months of age (**Fig. 2F**). Taken together, the *Snca*^*NLS/NLS*^ mice display age-dependent motor decline, gastrointestinal dysfunction, and premature lethality.

### *Snca*^*NLS/NLS*^ mice exhibit cortical atrophy, independent of αSyn aggregation or dopaminergic neurodegeneration

Many studies suggest that αSyn toxicity is intrinsically tied its aggregation, as the two are often associated in humans with PD and in animal models of the disease (1,30,31). However, models of αSyn toxicity often rely on the introduction of synthetically derived misfolded αSyn fibrils (32,33) or overexpression of αSyn (15,34), thereby potentiating its aggregation *in vivo*. Given that the *Snca*^*NLS*^ mice display age-dependent behavioral phenotypes, yet do not rely on αSyn overexpression, we asked whether accumulation of endogenous αSyn in the nucleus leads to its aggregation and thusly contributes to its toxicity. We examined both the solubility of αSyn as well as its pathologically-linked phosphorylation at serine residue 129 (pS129) by biochemical fractionation of brain samples of *Snca*^*NLS/NLS*^ mice compared to littermates. We used the *mThy1-SNCA* (“line 61”) transgenic and *Snca* knockout (*Snca*^*-/-*^) mouse lines as positive (34) and negative (35) controls, respectively. To our surprise, we found that the accumulation of nuclear αSyn does not lead to aggregation (**Fig. 3A**), nor does it become phosphorylated at S129 (**Fig. 3A, B**), and in fact total αSyn levels are reduced in these mice (**Fig. 3A-C**). These findings were further supported by histology, which showed no marked increase in pS129 in aged *Snca*^*NLS/NLS*^ mice compared to their respective littermates (**Fig. 3D**). This suggests that nuclear accumulation of αSyn confers toxicity independent of its aggregation.

**Figure 3:**
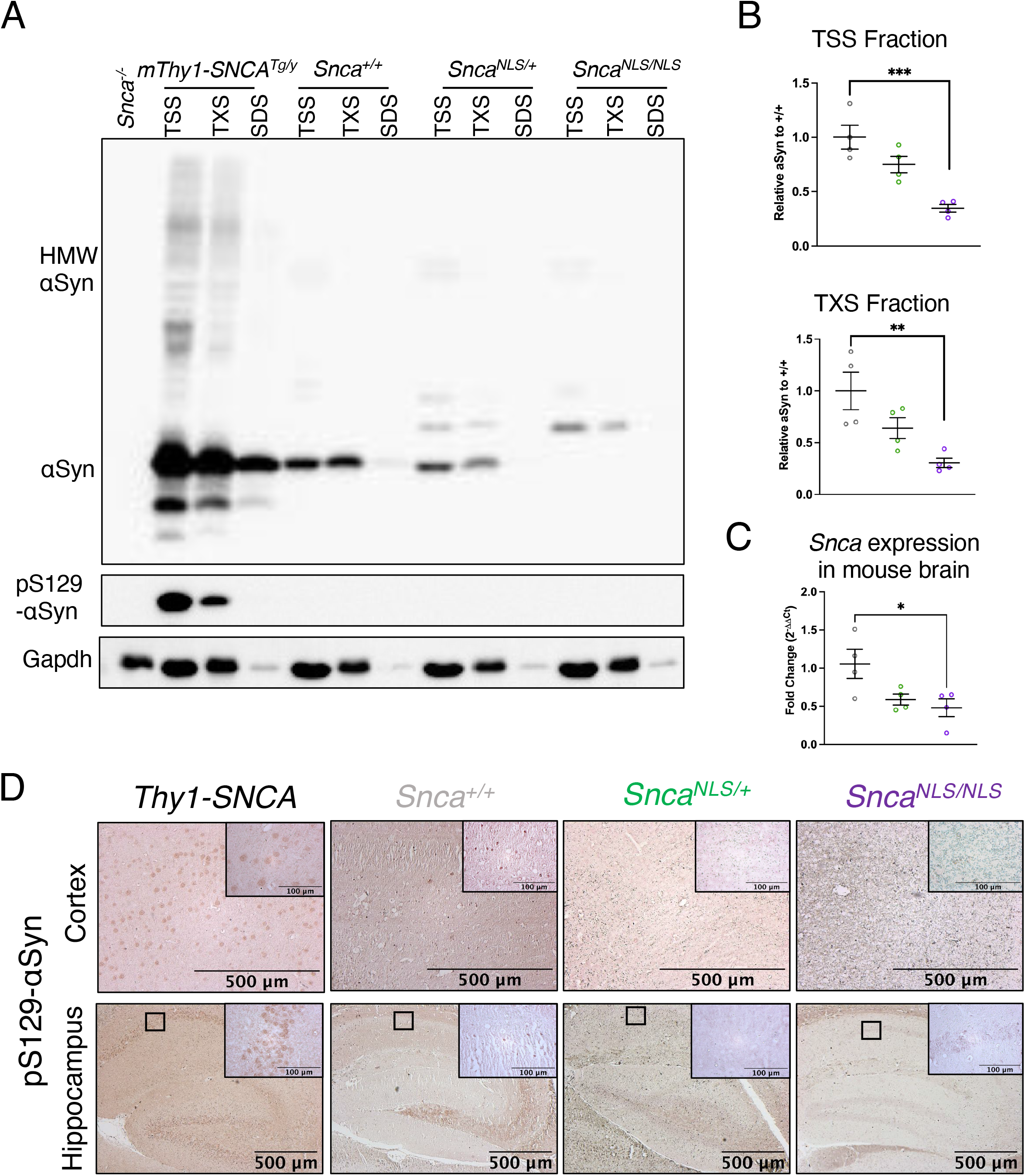
*Snca*^*NLS/NLS*^ mice do not display significant changes in aggregated or phosphorylated αSyn. Serial extraction of cortical mouse brain tissue with western blot probed for (A) αSyn (upper blot) and pS129-αSyn (middle blot) and Gapdh (bottom blot) comparing the level of αSyn in the (B) TSS (upper) and TXS (lower) fraction (n=4). C) qPCR of *Snca* mRNA from 9-month mouse cortex. D) pS129-αSyn staining of the motor cortex (upper) and hippocampus (lower). One-Way ANOVA with Bonferroni multiple comparison: ns, ^*, **^, and ^***^ denotes p>0.05, <0.05, <0.01, and <0.001, respectively.

A hallmark of PD is nigrostriatal dopaminergic neurodegeneration. Due to the relatively high expression of αSyn in dopaminergic neurons (**Fig. 1C**) (36), we hypothesized nuclear αSyn could be acutely toxic to dopaminergic neurons, causing their death, and ultimately leading to the observed behavioral deficits in *Snca*^*NLS*^ mice. To our surprise, we found that young and aged *Snca*^*NLS/NLS*^ mice had intact nigrostriatal tracts, when evaluated by striatal tyrosine hydroxylase (TH) fiber density and stereological estimation of dopaminergic cell number in the SNc at 3- and 18-months of age (**Fig. 4A-B, Supplementary Figure 3E**). Moreover, HPLC analysis of mouse striata revealed that 18-month-old mice across genotypes exhibit similar levels of dopamine and its metabolites (DOPAC, HVA, 5-HIAA; **Fig. 4C**).

**Figure 4:**
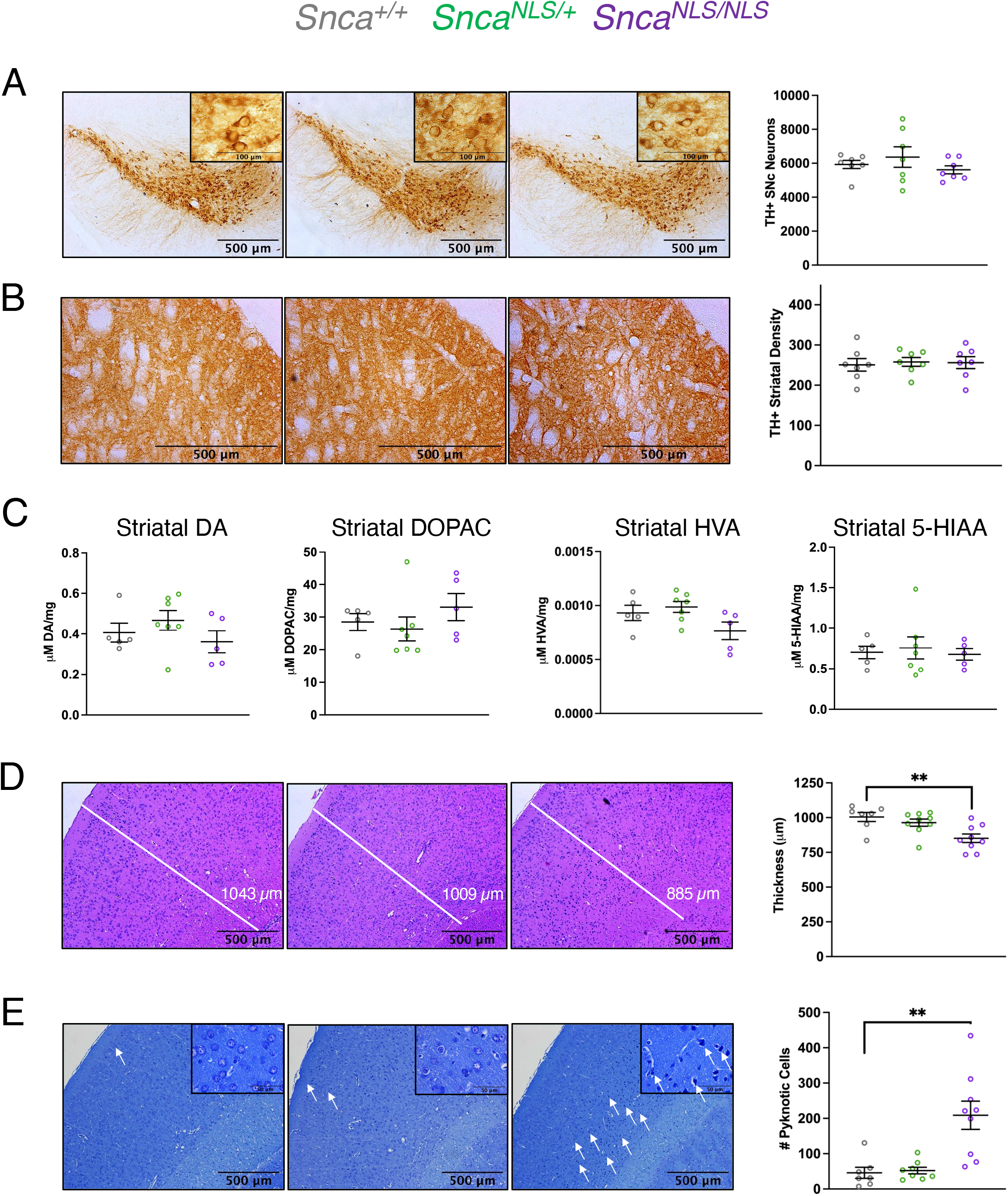
18-month-old *Snca*^*NLS/NLS*^ mice exhibit cortical thinning and pyknotic cells in the cortex independent of dopaminergic neurodegeneration. Tyrosine hydroxylase staining of the (A) Substantia nigra *pars compacta* and (B) striatum of 18-month mice (n=7). C) HPLC of 18-month striatal tissue measuring Dopamine (left), DOPAC (middle left), HVA (middle right), and 5-HIAA (right) (n=5-7). D) H&E staining with quantification of the motor cortex thickness (n=7-9). E) Toluidine blue staining with quantification of pyknotic cells from the motor cortex (n=7-9). White arrows denote select pykontic cells. One-Way with Bonferroni multiple comparison: ns, ^**^ denotes p>0.05, <0.01, respectively.

Since *Snca*^*NLS/NLS*^ mice exhibit motor defects without nigrostriatal degeneration nor αSyn aggregation, we took a step back to ask whether nuclear αSyn may impact other areas of the brain, thus contributing to PD-like phenotypes. Cortical involvement has long been linked to several synucleinopathies including PD, dementia with Lewy bodies (DLB), and PD with dementia (PDD) (31,37– 39). We therefore explored higher order cortical areas to determine whether *Snca*^*NLS/NLS*^ mice exhibit neurodegenerative features outside of the SNc. We conducted gross anatomical studies using hematoxylin and eosin (H&E) and toluidine blue staining and found significant anterior cortical thinning in the motor cortex (**Fig. 4D**) and a marked increase in pyknotic cells (**Fig. 4E**) throughout the cortex of 18-month-old *Snca*^*NLS/NLS*^ mice. These results suggest a potential etiology for the underlying motor behavior dysfunction in these mice.

### Unbiased proteomics uncovers reduced parvalbumin levels in *Snca*^*NLS/NLS*^ mice

We followed an unbiased strategy to uncover the molecular mechanisms underlying the behavioral and histological phenotypes of the *Snca*^*NLS/NLS*^ mice via quantitative proteomic analysis on cortices from 9-month-old mice. At this age, *Snca*^*NLS/NLS*^ mice exhibit robust behavioral abnormalities (**Fig. 2**), allowing identification of early molecular changes that drive the late-stage cortical atrophy exhibited in these mice and in PD. To quantify proteomic differences in wildtype and *Snca*^*NLS/NLS*^ mice, we performed pooled TMT10plex labeling for 5 wildtype and 5 *Snca*^*NLS/NLS*^ mouse cortices followed by mass spectrometry to identify differential proteomic changes (**Fig. 5A**). This approach yielded a list of nearly 1,800 proteins, of which 114 had a Log_2_ fold-change of ± 1 relative to wildtype (**Fig. 5B**). Of these 114 hits, 66 were downregulated and 48 were upregulated (**Supplementary Table 2**). GO term analysis revealed a significant enrichment for biological processes that are disturbed in PD, including regulation of GPCR signaling (**Fig. 5C**). To increase the stringency of our list, we filtered these hits using a statistical cut-off (Mann-Whitney *p*-value < 0.05). From this, we identified 10 high-confidence hits (**Fig. 5D**). Interestingly, among these 10 hits we noticed a few proteins of particular importance in DA signalling and have been associated with PD, such as Cacna1e, Darpp-32, Fgf1, Gng7, Pde10a, and SerpinA1a (40–47).

**Figure 5:**
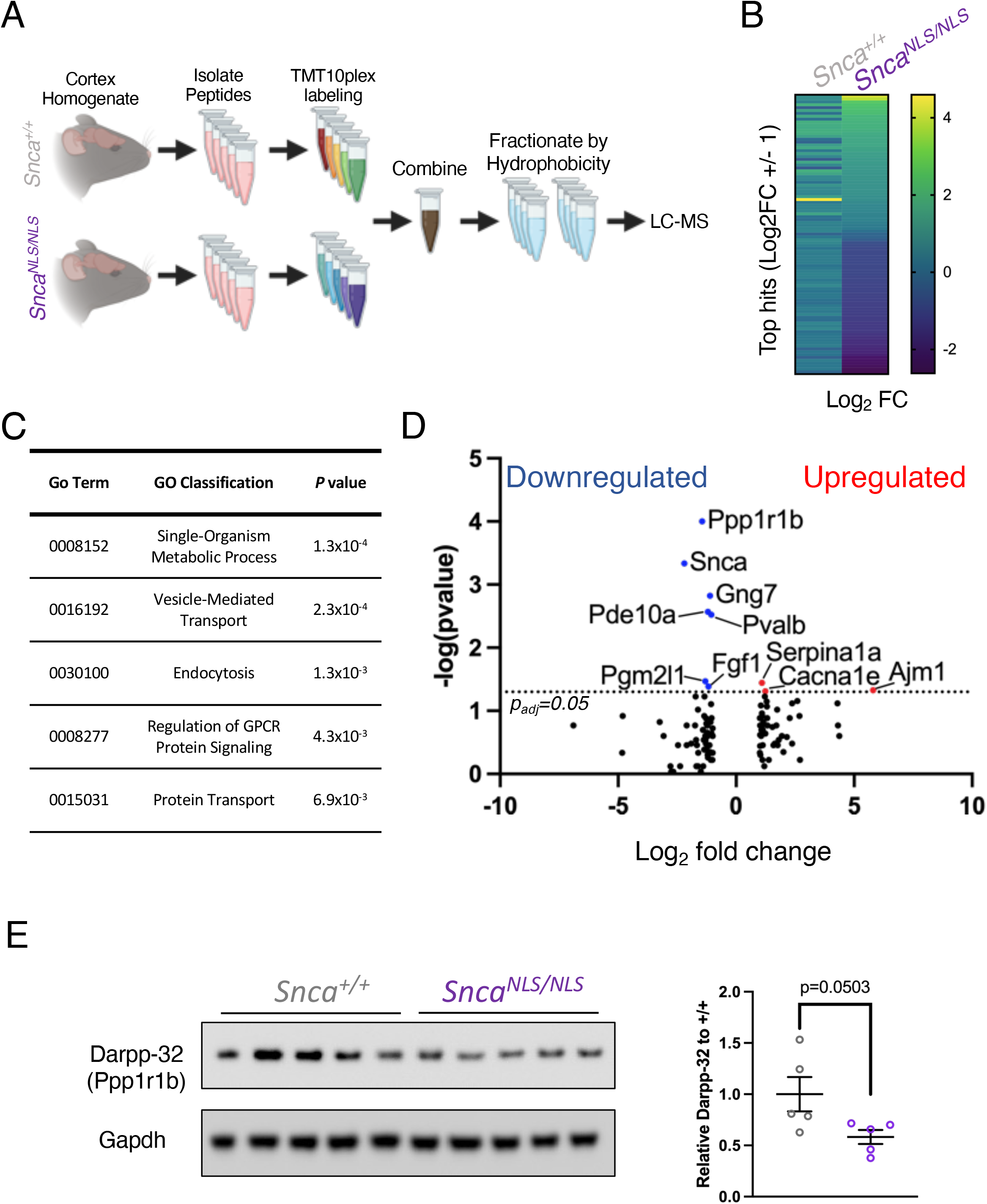
Mass Spectrometry reveals proteomic alterations in the *Snca*^*NLS/NLS*^ mice. A) Mass spectrometry scheme for quantitative comparison of proteome between 9-month wildtype and *Snca*^*NLS/NLS*^ mice (n=5). B) Heat map of all proteins identified through mass spectrometry with values within ±1 Log_2_ fold change. C) Table of enriched gene ontology pathways among the ± 1 Log_2_ fold change hits. D) Volcano plot of mass spectrometry results with ± 1 Log_2_ fold change highlighting significantly upregulated proteins in red and downregulated in blue. E) Western blot of cortical mouse brain tissue from 9-month mice probing for Darpp-32 (upper) and Gapdh (lower) with quantification (right). One-Way ANOVA with Bonferroni multiple comparison: ns denotes p>0.05.

Interestingly, we also observed a marked reduction in parvalbumin (PV), a protein recently shown to be disrupted in PD and in animal models of synucleinopathy (47). We confirmed the reduction of Darpp-32 in the *Snca*^*NLS/NLS*^ mice using western blot (**Fig. 5E**), thereby validating our proteomics approach. Collectively, these data suggest that disrupted dopaminergic signaling pathways that may underlie nuclear αSyn toxicity in PD.

## Discussion

The mechanisms underlying αSyn toxicity have been difficult to pin down. We and others have previously shown that nuclear αSyn is increased in the PD post-mortem brain and in animal models harboring *SNCA* mutations (6,13,20,26,48). Nonetheless, previous studies examining the role of nuclear αSyn in PD pathogenesis have yielded conflicting results, ranging from neurodegenerative (13,15,17,48) to neuroprotective (20,23) phenotypes. This may be in part due to the degree of overexpression of αSyn or the choice of read-out in these models. Our study sought to overcome this by answering if the nuclear accumulation of *native* αSyn is sufficient to cause PD-like phenotypes in mice. We engineered a mouse model with an *NLS-Flag* knockin on *Snca* to characterize the effects of chronically increased nuclear αSyn. We found that *Snca*^*NLS/NLS*^ mice reveal PD-like phenotypes including age-dependent motor decline and constipation, but also exhibit cortical atrophy, and reduced survival. The cortical atrophy we observed draws parallels to the cell loss seen in synucleinopathies with cortical involvement like DLB and PDD (31,37–39). Moreover, the anatomical location of this cell loss dovetails with the motor deficits seen in these mice and may shed light onto how nuclear αSyn in PD may be linked to cortical dysfunction and disease manifestation. When examining the proteomic profile of the *Snca*^*NLS/NLS*^ mice, we found a few high-confidence hits that have been previously associated with PD. Among these, we identified Pde10a, Darpp-32, SerpinA1a, parvalbumin, and Gng7. These are of particular interest as they hint at the potential role nuclear αSyn is playing in PD manifestation. Firstly, Pde10a and Darpp-32 are known to play a role in DA signalling, which may explain why the *Snca*^*NLS/NLS*^ mice exhibit a motor phenotype without loss in striatal DA. Secondly, a previous group also found a significant increase in SerpinA1a and decrease in Darpp-32 in mice injected with αSyn PFFs and transgenic mice that overexpress *hA53T-SNCA* (49).

Another PD mouse model, the *Thy1-SNCA* mice, shows loss of parvalbumin, which also occurs in PD patients (47,50). Additionally, we found a reduction in cortical Gng7. Interestingly, *Gng7* knockout mice exhibit a significant age-dependent motor deficits, particularly in the rotarod test, which was the motor assay with which the *Snca*^*NLS/NLS*^ mice had the most difficulty (51). Together, these changes in protein levels hint at the possible mechanisms whereby nuclear αSyn elicits toxicity.

While characterizing *Snca*^*NLS/NLS*^ mice, we consistently noted how this mouse line diverges from *Snca*^*-/-*^ mouse phenotypes (35), cementing that *Snca*^*NLS/NLS*^ mouse phenotypes are likely gain-of-function. To wit, *Snca*^*-/-*^ mice exhibit mild synaptic deficits in the absence of gross motor or non-motor deficits (35,52), likely due to compensation by β-synuclein and, to a lesser extent, γ-synuclein (53–55). Indeed, the motor phenotypes appear to be dependent on the local dose of nuclear αSyn as even the *Snca*^*NLS/+*^ mice exhibit some motor behavior deficits – albeit to a lesser extent than their *Snca*^*NLS/NLS*^ littermates, suggesting that they are due to a gain-of-function of nuclear αSyn and not a loss-of-function of synaptic αSyn. Nevertheless, we cannot exclude a model in which partial loss of synaptic αSyn combined with increased nuclear αSyn may drive the age-dependent behavioural and pathological phenotypes seen in *Snca*^*NLS*^ mice. Strikingly, behavioral and histological phenotypes in *Snca*^*NLS*^ mice occur independently from αSyn aggregation and pathogenic phosphorylation. This suggests a heretofore underappreciated role of *soluble*, nuclear αSyn in the pathogenesis of PD.

The cellular mechanisms that drive the nuclear accumulation of αSyn and its subsequent sequelae in PD remain elusive. Whether active or passive mechanisms bring αSyn to the nucleus is unknown. Native αSyn does not possess an NLS, therefore, it may be driven into the nucleus by passive mechanisms (it can traverse the nuclear pore complex due to its small size) (56) and could be kept there by interaction with nuclear components (e.g. histones or DNA) (16,17,20,22,23,57) or via uncharacterized modifications. Alternatively, active mechanisms such as its interaction with TRIM28 (26) or RAN (14) may be key in regulating its nuclear import. Moreover, αSyn likely has an important native role in the nucleus, particularly during mouse embryonic development, where nuclear αSyn constitutes up to 40% of its total cellular distribution, compared to 3-15% of total cellular distribution in adult mice (20). There, αSyn is suggested to bind both DNA and histones to modulate gene expression (16,17,22,23,57–60). In wildtype mice, nuclear αSyn was shown to be neuroprotective by binding to DNA and colocalizing with DNA damage response elements to protect against DNA damage (23). Whether the increase in nuclear αSyn observed in PD – modeled in the *Snca*^*NLS*^ mice – causes a gain of this normal developmental function or a neomorphic function will be important to establish, to facilitate future therapeutic development.

## Conclusion

This study tested whether a chronic accumulation of endogenous αSyn in the nucleus is sufficient to elicit PD-like phenotypes in mice. To do so, we generated a mouse allele with a nuclear localization signal on endogenous αSyn. These mice exhibit motor deficits, cortical atrophy, and a trend for reduced survival. Biochemical profiling of cortices from these mice identified changes in a handful of proteins involved in dopaminergic signaling, including Darpp-32, which may be key in driving these phenotypes. This new model enables the selective study of nuclear αSyn, thus allowing the field to amass a greater understanding of its role in disease independent of its often-studied aggregation.

## List of Abbreviations

5-HIAA: 5-hydroxyindoleacetic acid
αSyn: alpha-synuclein
DA: dopamine
Darpp-32: dopamine- and cAMP-regulated neuronal phosphoprotein
DOPAC: 3,4-dihydroxyphenylacetic acid
GO: Gene ontology
HPLC: high-performance liquid chromatography
HVA: homovanillic acid
NLS: nuclear localization signal
PD: Parkinson’s disease
PV: parvalbumin
SNc: Substantia Nigra *pars compacta*
TH: tyrosine hydroxylase

## Declarations

### Ethics approval

All animal work was performed under animal use protocols (breeding protocols CMMb-3009 and CMMb-3654, and experimental protocol CMMe-3091) approved by the University of Ottawa Animal Care Committee. The University of Ottawa is certified by the Canadian Council on Animal Care.

### Availability of data and materials

All data generated and analyzed during this study are included in this published article. The *Snca*^*NLS*^ mouse line will be made available through the Jackson Laboratory (Jax Stock No. 036763). Mass spectrometry data will be made available through Proteome Xchange.

### Authors’ contributions

H.M.G. generated the studied mouse cohorts, performed all behavioral tasks (with the help of K.M.R., Z.F., and S.M.C), and analyzed all figures included in this published article. K.M.R. also aided in tissue acquisition and processing and provided significant intellectual support. K.H. aided in the analysis of stained tissue. J.L.A.P. and T.R.S. harvested and maintained all primary cortical neurons. T.R.S. also obtained all primary neuron images and performed qPCR experiments. M.W.C.R conceptualized the study, designed the knockin mice, assisted in data collection and analysis. H.M.G. and M.W.C.R. wrote the manuscript and all authors provided edits.

## Acknowledgements

The authors thank M.G. Schlossmacher and J.J. Tomlinson (Ottawa Hospital Research Institute, OHRI) and M. Farrer (University of Florida) for *Snca*^*-/-*^ mice and R. Rissman and E. Masliah (University of California, San Diego) for the *mThy1-SNCA* (“line 61”) mice. The authors thank H.Y. Zoghbi and J.P. Revelli (Baylor College of Medicine) for initial project discussions, reagent development and the freedom to explore new ideas and bring mice to the Rousseaux lab with no strings attached. The authors also thank all members of the Rousseaux, Schlossmacher, and Zoghbi labs for important discussions and critical feedback on the manuscript. The authors thank K. Ure for her guidance in conceptualizing and analyzing behaviour tests. The authors also thank the Schlossmacher, M. Tiberi (OHRI), and S.X. Chen (uOttawa) labs for their generous donation of antibodies. The authors thank S.X. Chen and K. Ure for providing critical feedback to the manuscript. The authors also thank the following Core facilities from the University of Ottawa and the Ottawa Hospital Research Institute for use of their facility, equipment, and expertise: Animal Behaviour and Physiology Core, Cell Biology and Imaging Acquisition Core, Louise Pelletier Histology Core, and the OHRI Proteomics Core. The authors also thank the Genome Engineered Rodent Models Core at Baylor College of Medicine and Animal Care and Veterinary Service. Figs. 1D, 5A, and Supplementary Figures 1A,C were generated in part with Biorender.com. All histograms were generated with Prism 9.

## Figures, tables, additional files

**Supplementary Figure 1:**
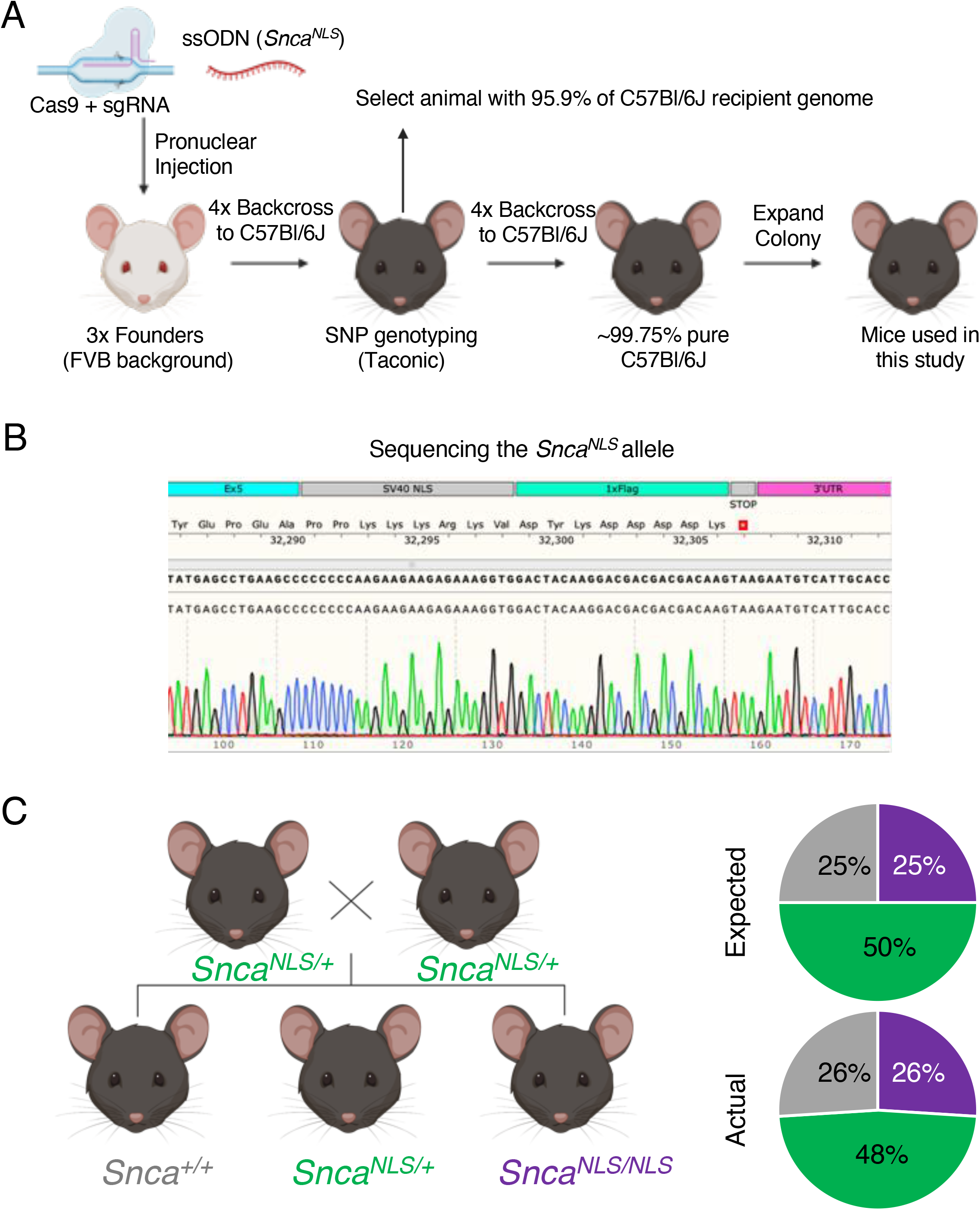
Generation of the *Snca*^*NLS-Flag*^ mice. A) *Snca*^*NLS-Flag*^ mouse generation. B) Sequencing confirms the presence of the knock-in. C) Breeding scheme with expected and actual Mendelian ratios among offspring. n=148 *Snca*^*+/+*^, n=272 *Snca*^*NLS/+*^, and n=147 *Snca*^*NLS/NLS*^.

**Supplementary Figure 2:**
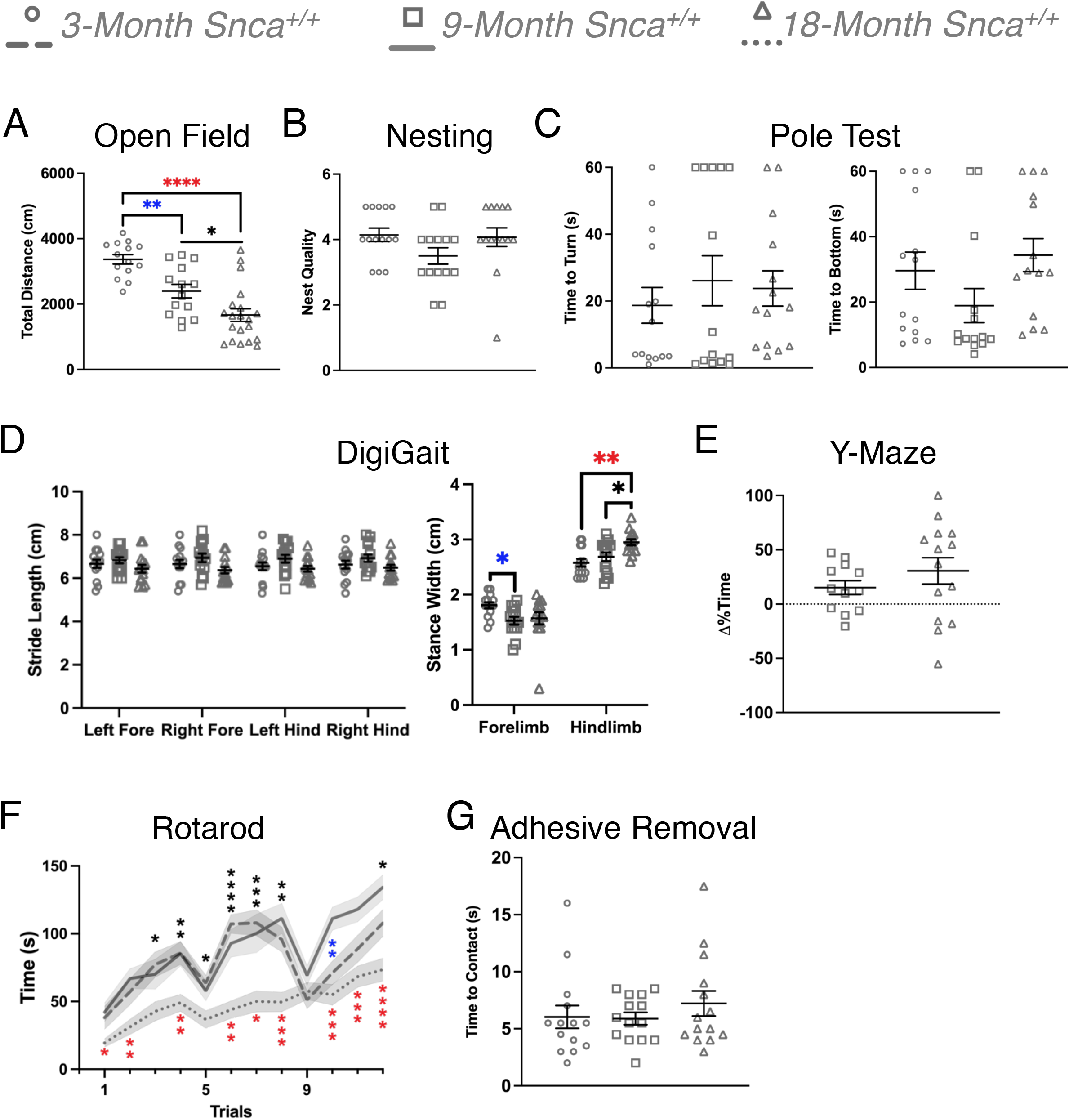
Wild-type mice exhibit age-dependent decline in motor ability. Behavior comparison of wild-type mice at 3-, 9-, and 18-months in (A) Open Field, (B) Nesting, (C) Pole test measuring time to turn over (left) and time to descend to the bottom (right), (D) Digigait measuring stride length (left) and stance width (right), and (E) Y maze (n=14-20). Motor assays with additional cohorts of mice include (F) Rotarod and (G) Adhesive removal measuring time to contact their forepaws (n=9-20). One-(A,B,C,E,G) or Two-Way ANOVA (D,F): ns, ^*, **, ***^, and ^****^ denotes p>0.05, <0.05, <0.01, <0.001, and <0.0001, respectively. Blue asterisk denotes significance between 3- and 9-month mice, black asterisk denotes significance between 9- and 18-month mice, and red asterisk denotes significance between 3- and 18-month mice.

**Supplementary Figure 3:**
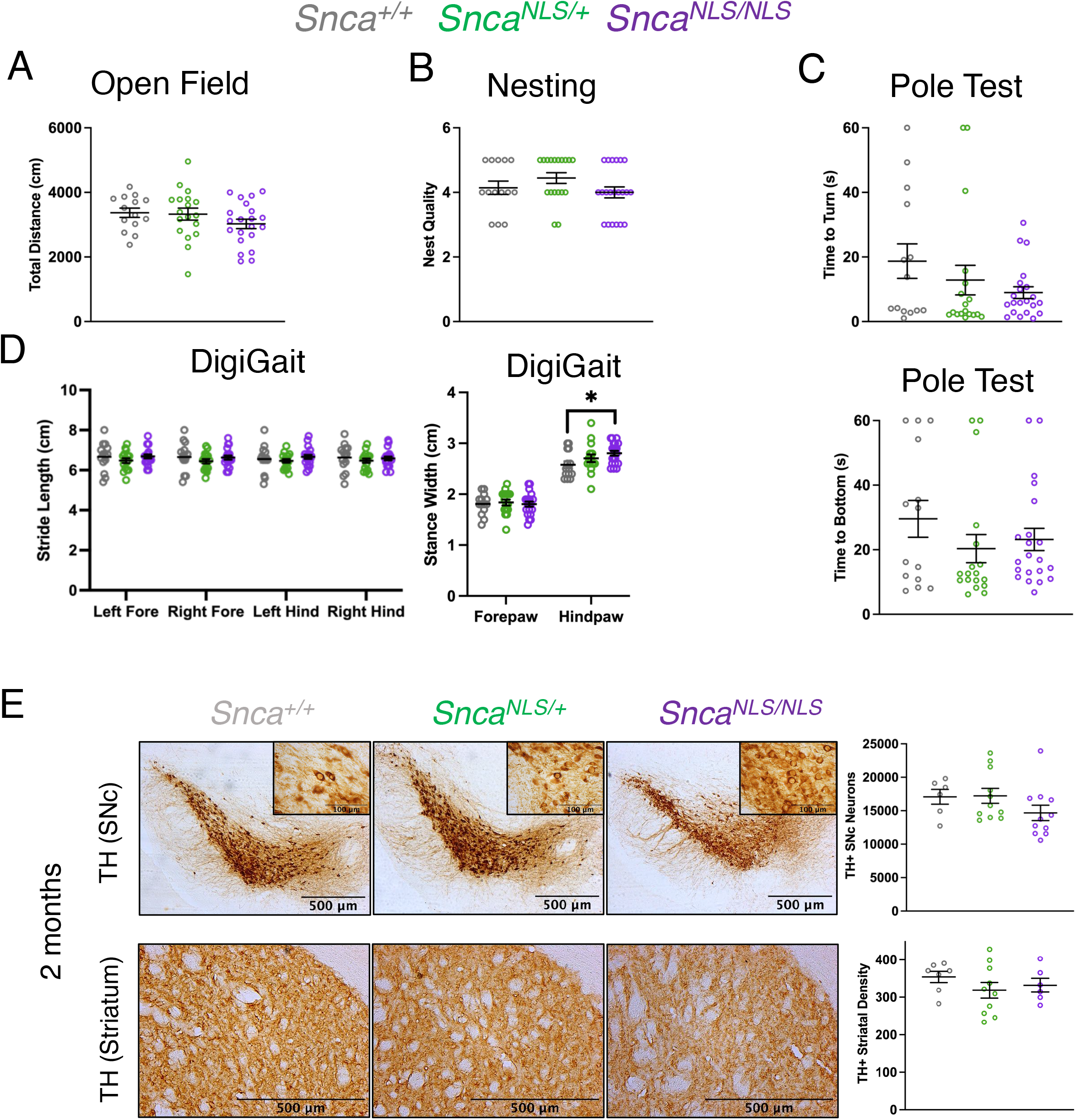
Comprehensive behavior and dopaminergic profiling of young *Snca*^*NLS/NLS*^ mice reveals little-to-no phenotypes. Behavior analysis of 3-month mice in (A) Open Field, (B) Nesting, (C) Pole test measuring time to turn over (upper) and time to descend to the bottom (lower), and (D) DigiGait measuring stride length (left) and stance width (right) (n=14-21). E) Tyrosine hydroxylase staining of the Substantia nigra *pars compacta* (upper) and striatum (lower) of 2-month mice with their respective quantifications (right) (n=6-11). One-Way ANOVA (A,B,C,E) or Two-Way ANOVA (D): ns, ^*^ denotes p>0.05, <0.05, respectively.

**Supplementary Figure 4:**
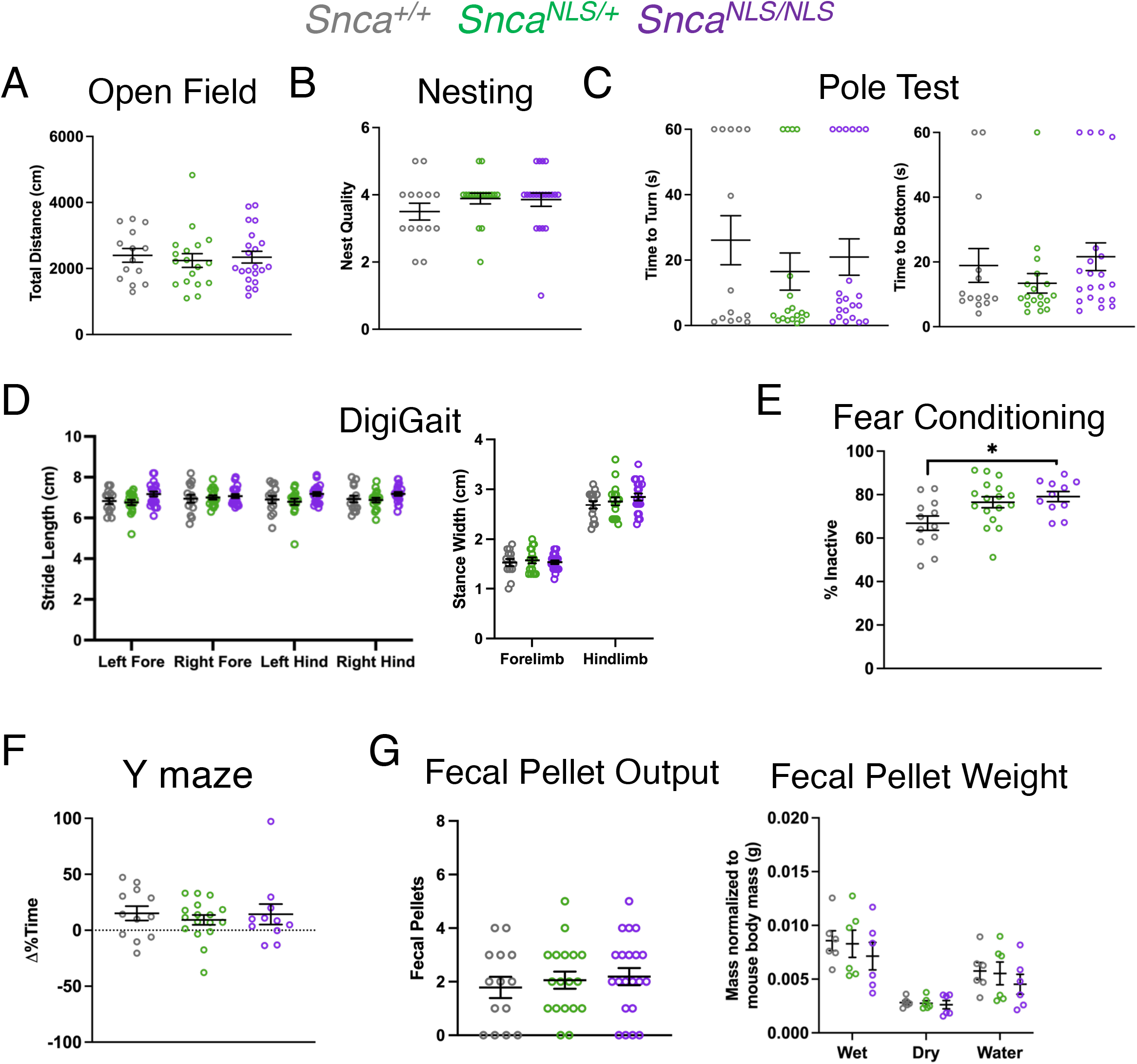
Comprehensive behavior profiling of 9-months-old *Snca*^*NLS/NLS*^ mice compared to littermates. Behavior analysis of 9-month mice in motor assays including (A) Open Field, (B) Nesting, (C) Pole test measuring time to turn over (left) and time to descend to the bottom (right), and (D) DigiGait measuring stride length (left) and stance width (right). Non-motor assays include (E) Fear conditioning, (F) Y maze forced alteration, and (G) Fecal pellet production measuring fecal output in 10 minutes (left) and the weight and water content of fecal pellets produced over 1 hour (right) (n=14-21). One-Way ANOVA (A,B,C,E,F,G left) or Two-Way ANOVA (D,G right): ns, ^*^ denotes p>0.05, <0.05, respectively.

**Supplementary Figure 5:**
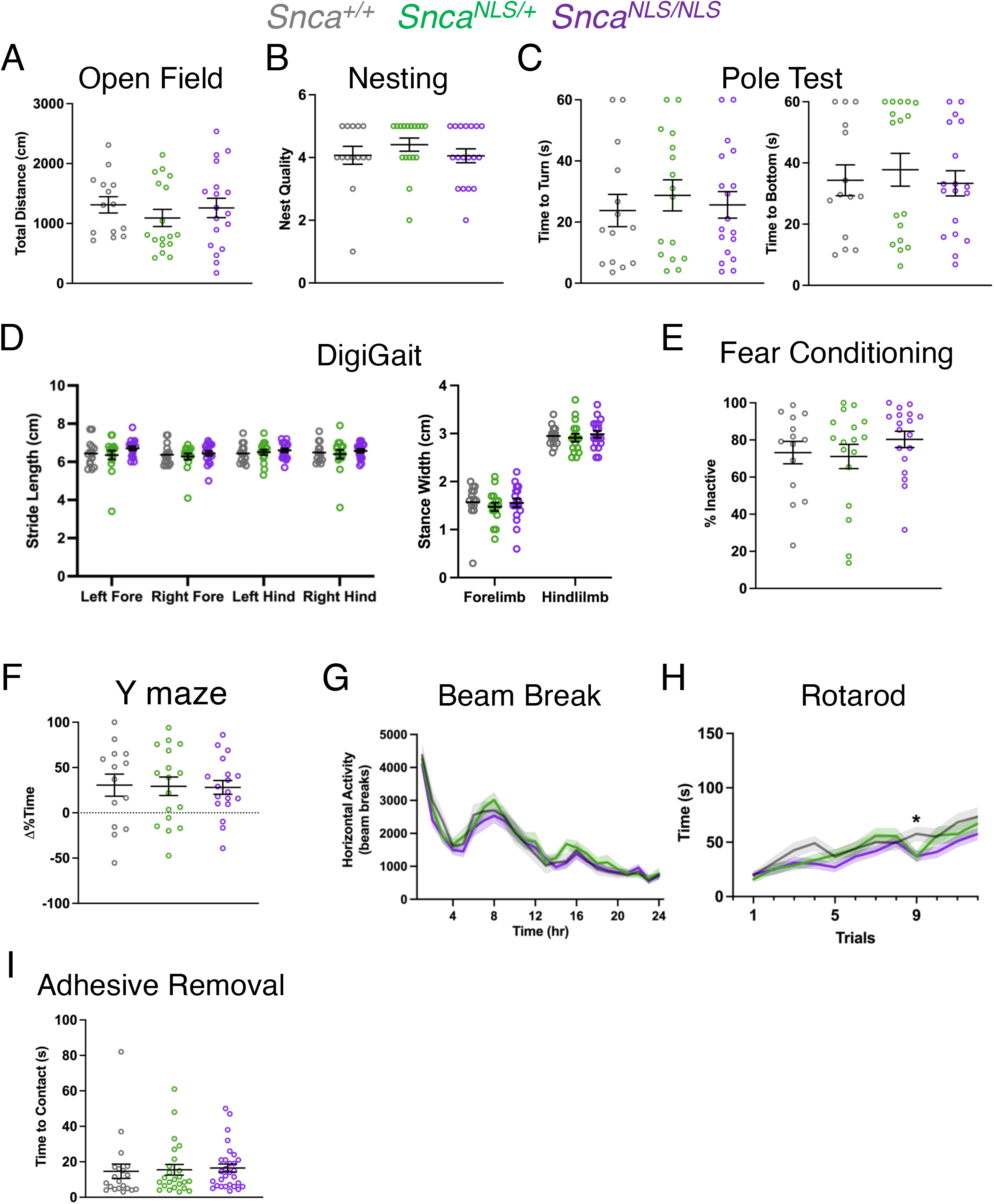
Comprehensive behavior profiling of 18-months-old *Snca*^*NLS/NLS*^ mice compared to littermates. Behavior analysis of 18-month mice in motor assays including (A) Open Field, (B) Nesting, (C) Pole test measuring time to turn over (left) and time to descend to the bottom (right), and (D) Digigait measuring stride length (left) and stance width (right). Non-motor assays include (E) Fear conditioning and (F) Y maze (n=14-18). Motor assays with additional cohorts of mice include (G) Beam Break, (H) Rotarod, and (I) Adhesive removal measuring time to contact their forepaws (n=20-31). One-Way ANOVA (A,B,C,E,F,I) or Two-Way ANOVA (D,G,H): ns, ^*^ denotes p>0.05, <0.05, respectively.

**Supplementary Table 1: List of antibodies used throughout the study**

**Supplementary Table 2: List of hits from proteomic profiling of cortices from 9-month-old *Snca***^***NLS/NLS***^ **mice compared to wild-type littermates**

